# Automated discovery of relationships, models and principles in ecology

**DOI:** 10.1101/027839

**Authors:** Pedro Cardoso, Paulo A. V. Borges, José C. Carvalho, François Rigal, Rosalina Gabriel, José Cascalho, Luís Correia

## Abstract

1. Ecological systems are the quintessential complex systems, involving numerous high-order interactions and non-linear relationships. The most commonly used statistical modelling techniques can hardly reflect the complexity of ecological patterns and processes. Finding hidden relationships in complex data is now possible through the use of massive computational power, particularly by means of Artificial Intelligence methods, such as evolutionary computation.
2. Here we use symbolic regression (SR), which searches for both the formal structure of equations and the fitting parameters simultaneously, hence providing the required flexibility to characterize complex ecological systems.
3. First, we demonstrate how SR can deal with complex datasets for: 1) modelling species richness; and 2) modelling species spatial distributions. Second, we illustrate how SR can be used to find general models in ecology, by using it to: 3) develop species richness estimators; and 4) develop the species-area relationship and the general dynamic model of oceanic island biogeography.
4. All the examples suggest that evolving free-form equations purely from data, often without prior human inference or hypotheses, may represent a very powerful tool for ecologists and biogeographers to become aware of hidden relationships and suggest general theoretical models and principles.

## INTRODUCTION

### Ecology as a complexity science

Complexity is a term often used to characterize systems with numerous components interacting in ways such that their collective behaviour is difficult to predict, but where emergent properties give rise to, more or less simple but seldom linear, patterns (Table 1)(Holland 1995; Mitchell 2009). Complexity science is therefore an effort to understand non-linear systems with multiple connected components and how “the whole is more than the sum of the parts” (Holland 1998). Biological systems probably are among the most complex (Solé & Goodwin 2000), and among them, ecological systems are the quintessential complex systems (Anand 2010). These are composed of individuals, populations from different species, interacting and exchanging energy in multiple ways, furthermore relating with the physical environment at different spatial and temporal scales in non-linear relationships. As a consequence, ecology is dominated by idiosyncratic results, with most ecological processes being contingent on the spatial and temporal scales in which they operate, which makes it difficult to identify recurrent patterns, knowing also that pattern does not necessarily identify process (Lawton 1996; Dodds 2009; Passy 2012). The most commonly used exploratory (e.g. PCA, NMDS) and statistical modelling techniques (e.g.linear and non-linear regression) can hardly reflect the complexity of ecological patterns and processes, often failing to find meaningful relationships in data. More flexible techniques (e.g. GAMs) usually do not allow an easy interpretation of results and particularly of putative causal relationships. For ecological data, we require more flexible and robust, yet amenable to full interpretation, analytical methods, which can eventually lead to the discovery of general principles and models.

**Table 1.**
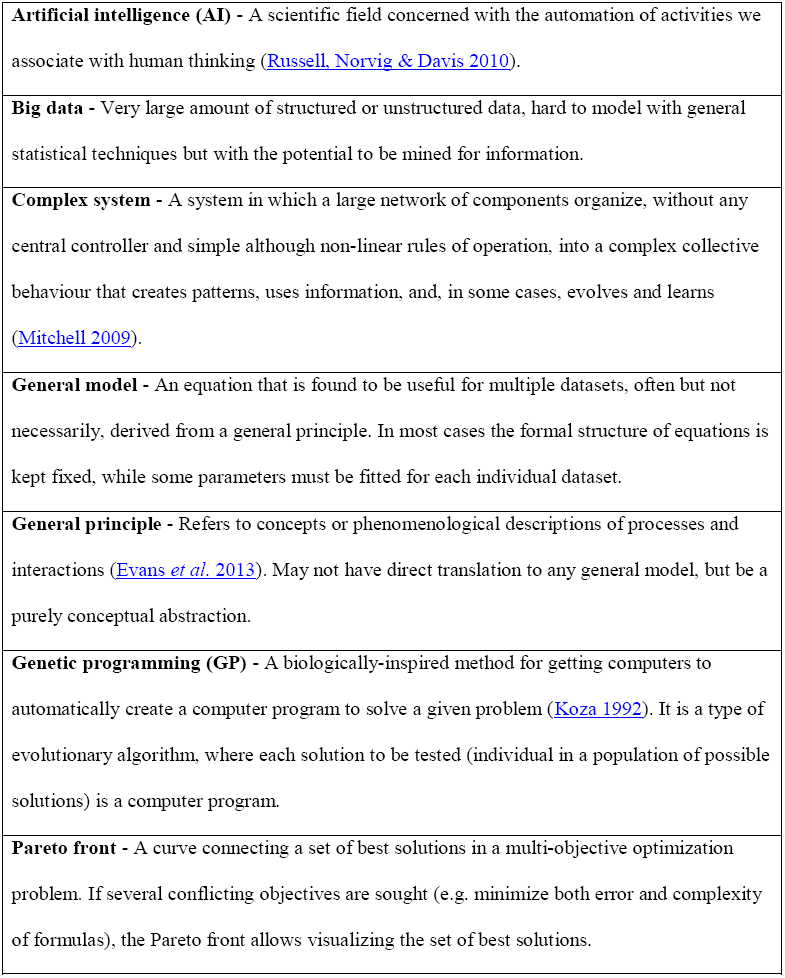
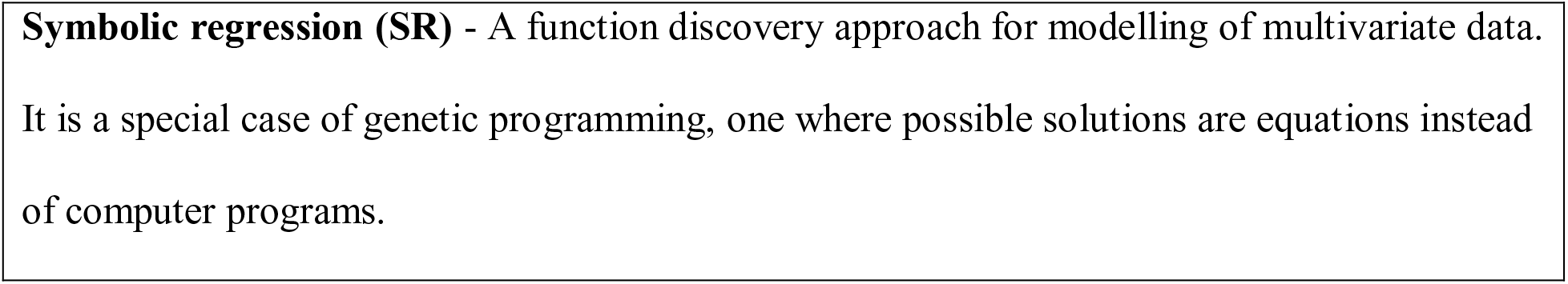
Glossary of Terms.

### General principles and models in ecology

The ultimate aim of any ecological principle is to provide a robust model for exploring, describing and predicting ecological patterns and processes regardless of taxon identity and geographic region (Lawton 1996; Dodds 2009). Finding a recurrently high goodness-of-fit for a model to an ecological pattern for most taxa and ecosystems is usually the most compelling evidence of a mechanistic process controlling that pattern. When general principles are translated into robust models, general statistical methods are mostly abandoned in favor of these (Appendix 1). Such general, widely applicable, equations are mostly found by intellectual *tour de force*. Yet, they surely are only the tip of the iceberg, usually incorporating few of the variables increasingly available to ecologists and that could potentially explain such patterns.

### Computing power applied to complex ecological systems

The automation of techniques for collecting and storing ecological and related data, with increasing spatial and temporal resolutions, has become one of the central themes in ecology and bioinformatics. Yet, automated and flexible ways to synthesise such complex and big data were mostly lacking until recently. Finding hidden relations within such data is now possible through the use of massive computational power. New computer-intensive methods have been developed or are now available or possible (Reshef *et al.* 2011) including in particular the broad field of Artificial Intelligence (AI) which has produced a variety of approaches. AI includes a series of evolution-inspired techniques, brought together in the sub-field of evolutionary computation, of which the most studied and well-known probably are genetic algorithms (Holland 1975). Genetic programming, namely in the form of symbolic regression (SR)(Koza 1992), is a particular derivation of genetic algorithms that searches the space of mathematical equations without any constraints on their form, hence providing the required flexibility to represent complex systems as presented by many ecological systems (Fig 1). Contrary to traditional statistical techniques, symbolic regression searches for both the formal structure of equations and the fitting parameters simultaneously (Schmidt & Lipson 2009). Finding the structure of equations is especially useful to discover general models, providing general insights into the processes and eventually leading to the discovery of new and as yet undiscovered principles. Fitting the parameters provides insight into the specific data, and allow specific predictions.

**Figure 1.**
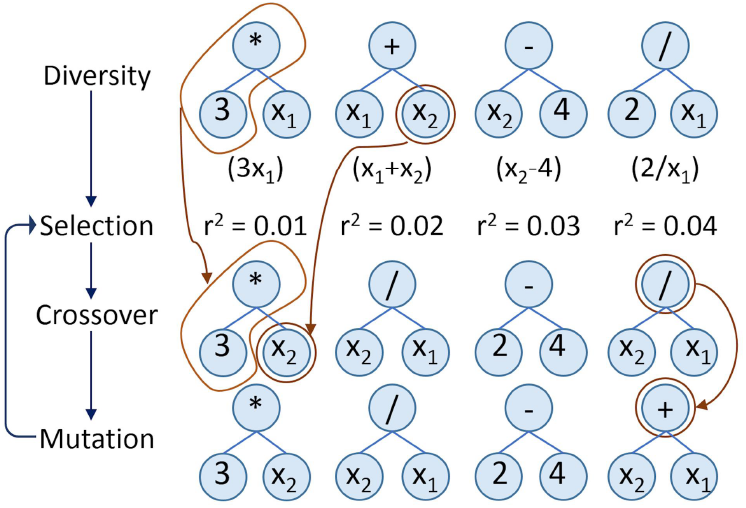
Schematic representation of the symbolic regression workflow. The basic representation is a parse-tree where building blocks such as variables (in this case: x_1_, x_2_), parameters (integers or real numbers) and operators (e.g. +, −, ×, ÷) are connected forming functions (in parenthesis under the first line of trees). Initial equations are generated by randomly linking different building blocks. Equations are combined through crossover, giving rise to new equations with characteristics from both parents (arrows linking the first and second rows of trees). Equations with better fitness (e.g. r^2^) have higher probabilities of recombining. To avoid loss of variability, a mutation step is added after crossover (arrows linking the second and third rows of trees). After multiple generations, evolution stops and a set of free-form equations best reflecting the input data is found.

Successful examples on the use of SR in ecology include modelling of land-use change (Manson 2005; Manson & Evans 2007), effects of climate change on populations (Tung *et al.* 2009; Larsen *et al.* 2014), community distribution (Larsen, Field & Gilbert 2012; Yao *et al.* 2014), predicting micro-organismal blooms (Muttil & Lee 2005; Jagupilla *et al.* 2015), deriving vegetation indices (Almeida *et al.* 2015) and using parasites as biological tags (Barrett, Kostadinova & Raga 2005).

In this work we explain, test and demonstrate the usefulness of SR in uncovering hidden relationships within typical ecological datasets. First, we demonstrate how SR can deal with complex datasets, namely for: 1) modelling species richness; and 2) modelling species spatial distributions. Second, we illustrate how SR can be used to find general models in ecology, by using it to: 3) develop species richness estimators; and 4) develop the species-area relationship (SAR) and the general dynamic model of oceanic island biogeography (GDM).

## Materials and Methods

### Symbolic regression

Symbolic regression works as a computational parallel to the evolution of species (Fig 1). A population of initial equations is generated randomly by combining different building blocks, such as the variables of interest (independent explanatory variables), algebraic operators (+, −, ÷, ×), analytic function types (exponential, log, power, etc.), constants and other ways to combine the data (e.g. Boolean or decision operators). Being random, these initial equations almost invariably fail, but some are slightly better than others. All are then combined through crossover, giving rise to new equations with characteristics from both parents. Equations with better fitness (e.g. higher r^2^) have a higher probability of recombining. To avoid new equations being bounded by initially selected building blocks or quickly losing variability along the evolutionary process, a mutation step (acting on any building block) is added to the process after crossover. After multiple generations, an acceptable level of accuracy by some of the equations is often attained and the researcher stops the process.

For this work we used the software Eureqa (Schmidt 2015). For each run, the software outputs a list of equations along an error/complexity Pareto front, with the most accurate equation for each level of complexity being shown (Fig 2). The Pareto front often presents an “elbow”, where near-minimum error meets near-minimum complexity. The equation in this inflection is closer to the origin of both axes and is a good starting point for further investigation – if both axes are in comparable qualitative scales. Often, however, this inflection point is not obvious and a single formula is not clearly best. In such cases, weights can be given to each of them through Bayesian statistics, using indices that positively weight accuracy and negatively weight complexity, such as Akaike’s Information Criterion - AIC (Akaike 1974). However, in all cases it is important to check all formulas along the Pareto front. Often equations or models that make immediate sense to the specific question may not be detected by these automated methods.

**Figure 2.**
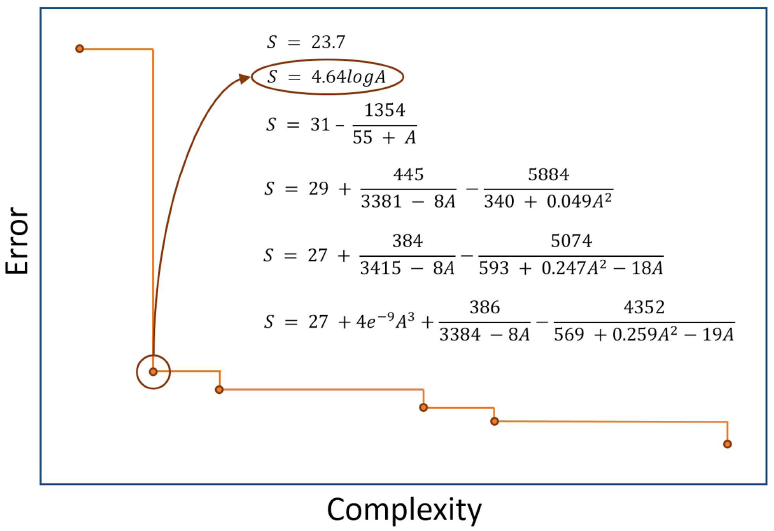
Example of a Pareto front depicting error vs. complexity. This example reflects a symbolic regression search of the best species–area relationship for native spiders in the Azores (Portugal). The second formula is clearly the most promising, with both high accuracy (low error) and low complexity. In many occasions a single formula is not clearly best, in which case weights can be given to each of them through Bayesian statistics and multiple formulas presenteds possible outcomes.

### Case-studies

#### Modelling species richness

Modelling and mapping the species richness of high diversity taxa at regional to large scales is often impossible without extrapolation from sampled to non-sampled sites. Here, we used an endemic arthropod dataset collected in Terceira Island, Azores. Fifty-two sites were sampled using pitfall traps for epigean arthropods (Cardoso *et al.* 2009), 13 in each of four land-use types: natural forest, exotic forest, semi-natural pasture and intensively managed pasture. This dataset was randomly divided into training and test data (26 sites each). We explained and predicted species richness per site using elevation, slope, annual average temperature, annual precipitation and an index of disturbance (Cardoso *et al.* 2013).

As the response variable was count data, Generalized Linear Models (GLM) and Generalized Additive Models (GAM) with a Poisson error structure with log link were used. We used the package MuMIn (Barton 2015) and the R environment (Team 2015) for multi-model inference based on AICc values, using all variables plus all possible interactions for GLM. For GAM we used package gam (Hastie 2015). For the SR search we used only algebraic and analytic operators (+, −, ÷, ×, log, power), in this and all examples below, so that outputs could be most easily interpreted. The r^2^ goodness of fit was used as the fitness measure. As there was no clearly best formula, AICc was used to choose a single equation along the Pareto front (Appendix 2). Both r^2^ and AICc were used to compare GLM and GAM with SR on the test dataset. Here and in subsequent analyses, all models with a ΔAICc value < 2 (the difference between each model’s AICc and the lowest AICc) were considered as receiving equal statistical support.

#### Modelling species distributions

Species distribution modelling (SDM) is widely used to fill gaps in our knowledge on individual species distributions. One of the general statistical methods used for SDM is logistic regression. Among the multiple alternatives, the principle of maximum entropy (Maxent)(Phillips, Anderson & Schapire 2006) has been found to be particularly robust (Elith *et al.* 2006).

We modelled the potential distribution of two endemic Azorean species in Terceira Island: the rare forest click-beetle *Alestrus dolosus* (Coleoptera, Elateridae) and the abundant but mostly forest restricted spider *Canariphantes acoreensis* (Araneae, Linyphiidae). Given the intrinsic differences between methods, we had to use different background datasets. Maxent used the environmental maps of the islands with a resolution of 100 m, from where it extracted pseudo-absences. We then converted the probabilistic potential distribution maps to presence/absence using the maximum value of training sensitivity plus specificity as the threshold as recommended by Liu et al. (Liu *et al.* 2005). Logistic regression and SR used presence/absence data from the 52 sampled sites. We used the package MuMIn (Barton 2015) and the R environment (Team 2015) for multi-model inference of logistic regression based on AICc values. In the SR run a step function was included, so that positive and negative values were converted to presence and absence, respectively. Absolute error, reflecting the number of incorrect classifications, was used as the fitness measure. As inflection points of the Pareto fronts were clear, the best SR formula for each species was chosen based on them (Appendix 2). In all cases only the training data (26 sites) were used for running the models. Logistic regression, Maxent and SR were compared in their performance for predicting presence and absence of species on the 26 test sites using the True Skill Statistic - TSS (Allouche, Tsoar & Kadmon 2006).

#### Developing species richness estimators

Several asymptotic functions have been used to estimate species richness (Soberon & Llorente 1993), including the Clench function (Clench 1979), the negative exponential function and the rational function (Ratkowsky 1990) (Appendix 1). Our objective was to rediscover or eventually find asymptotic models that would outperform them. Two independent datasets were used resulting from exhaustive sampling for spiders in 1ha plots, performed by 8 collectors during 320 hours of sampling in a single hectare using five different methods. The training dataset was from a mixed forest in Gerês (northern Portugal) and the test dataset was from a *Quercus* forest in Arrábida (southern Portugal) (Cardoso *et al.* 2008a; Cardoso *et al.* 2008b).

Randomized accumulation curves for both sites were produced using the R package BAT (Cardoso, Rigal & Carvalho 2015) (the package also includes both datasets). The true diversity of each site was calculated as the average between different non-parametric estimators (Chao 1 and 2, Jackknife 1 and 2). Because the sampled diversity in the training dataset reached a very high completeness but we wanted to simulate typically very incomplete sampling, datasets with 10, 20, 40, 80 and 160 randomly chosen samples were extracted and used, in addition to the complete 320 samples dataset, as independent runs in SR. Squared error was used as the fitness measure. Additionally, we imposed a strong penalty to non-asymptotic functions, although these were still allowed in the search process. The weighted and non-weighted scaled mean squared errors implemented in BAT (Cardoso, Rigal & Carvalho 2015) were used as accuracy measures.

#### Developing the species-area relationship (SAR) and the general dynamic model of oceanic island biogeography (GDM)

One of the most studied examples of SARs is their application to island biogeography (ISAR). The shape of ISARs has been modelled by many functions, but three of the simplest seem to be preferred in most cases, the power, exponential and linear models (Triantis, Guilhaumon & Whittaker 2012)(Appendix 1).

The general dynamic model of oceanic island biogeography was proposed to account for diversity patterns within and across oceanic archipelagos as a function of area and age of the islands (Whittaker, Triantis & Ladle 2008). Several different equations have been found to describe the GDM, extending the different SAR models with the addition of a polynomial term using island age and its square (TT^2^), depicting the island’s ontogeny. The first to be proposed was an extension of the exponential model (Appendix 1)(Whittaker, Triantis & Ladle 2008), the power model extensions following shortly after (Fattorini 2009; Steinbauer 2013).

Our objective was to test if we could re-discover and eventually refine existing models for the ISAR and GDM from data alone. We used the Azores and Canary Islands spiders (Appendix 3)(Cardoso *et al.* 2010) as training data. To independently test the generality of models arising from spider data, we used bryophyte data from the same archipelagos (Appendix 3)(Aranda *et al.* 2014). The area and maximum time since emergence of each island were used as explanatory variables and the native species richness per island as the response variables. The r^2^ value was used as the fitness measure. The best SAR and GDM equations found by SR were chosen based on the inspection of the Pareto front (Appendix 2), but looking also for interpretability of the models. These were then compared with the existing models using AICc using the R package BAT (Cardoso, Rigal & Carvalho 2015).

## RESULTS

### Modelling species richness

The model selected by GLM was:

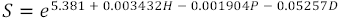

(r^2^ = 0.744, AICc = 30.793), where H = altitude, P = precipitation and D = disturbance. Yet, the GLM model seems to be overfitting, as the results with the test data were considerably worse (r^2^ = 0.146, AICc = 63.672). Overfitting also occurred with GAM, as the model was extremely good for the training data (r^2^ = 0.930, AICc = 8.643) yet much worse for testing data (r^2^ = −0.077, AICc = 85.601). The SR results performed worse than GLM or GAM with the training data, with the formula chosen according to AICc being:

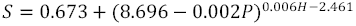

(r^2^ = 0.641, AICc = 43.050). However, the SR equation performs considerably better than GLM or GAM with the test data (r^2^ = 0.289, AICc = 62.354), revealing a higher generality of this formula.

### Modelling species distributions

The potential distribution models are relatively similar for *C. acoreensis* but show marked differences for *A. dolosus* (Fig 3). Symbolic regression outperforms both other models for *A. dolosus* and is as good as Maxent for *C. acoreensis*, with both outperforming LR (Table 2). The SR models are not only the best, presenting maximum values for TSS, but are also the easiest to interpret. *A. dolosus* is predicted to have adequate environmental conditions in all areas above 614m elevation, being restricted to pristine native forest. *C. acoreensis* can potentially be present in all areas with disturbance values below 41.3, occurring not only in native forest but also in adjacent semi-natural grassland and humid exotic forest. The LR and Maxent models used a large number of explanatory variables for *A. dolosus*, yet performed worse on the test data than did SR.

**Figure 3.**
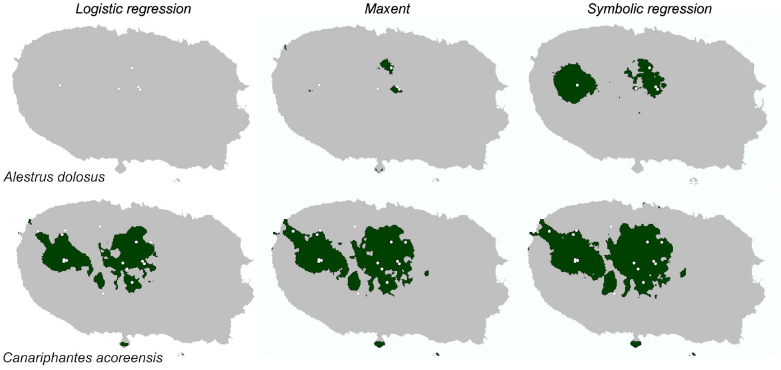
Predicted distribution of two Azorean arthropods using three modelling methods. Observed locations (white dots) and predicted distribution (dark green areas) of *Alestrus dolosus* (Coleoptera) and *Canariphantes acoreensis* (Araneae) in the island of Terceira (Azores, Portugal) using logistic regression, maximum entropy and symbolic regression.

**Table 2.** Species distribution models for two endemic arthropod species on the island and of Terceira (Azores, Portugal). Accuracy statistics on an independent test dataset are given by the True Skill Statistic (TSS). *H* =altitude, *Sl* = slope, *T* = average annual temperature, *P* = annual precipitation and *D* =disturbance index. The step function in symbolic regression converts positive values inside parentheses to presence and negative values to absence. Best values in bold.

| Model | Formula | Sensitivity | Specificity | TSS |
| --- | --- | --- | --- | --- |
| *Alestrus dolosus* |  |  |  |  |
| Logistic regression | $\frac{1}{1 + e^{-(8469 - 0.432P - 540.7T)}}$ | 0 | **1** | 0 |
| Maxent | Uses all variables but *Sl*, main is *D*<br>(contribution = 74.1%) | 0.5 | **1** | 0.5 |
| Symbolic regression | $step(H - 614)$ | **1** | 0.75 | **0.75** |
| *Canariphantes acoreensis* |  |  |  |  |
| Logistic regression | $\frac{1}{1 + e^{-(3.617 - 0.103D)}}$ | 0.667 | **0.7** | 0.367 |
| Maxent | Uses only *D* (contribution = 100%) | **0.833** | 0.65 | **0.483** |
| Symbolic regression | $step(41.3 - D)$ | **0.833** | 0.65 | **0.483** |

### Developing species richness estimators

For the training dataset, one asymptotic model was found by SR (Appendix 2):

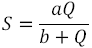

where *a* and *b* are fitting parameters. This model is in fact the Clench model with a different formulation (Appendix 1), where the asymptote is *a.* A second, slightly more complex but better fitting, model was found for partial datasets with 40 or more samples:

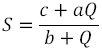

where *c* is a third fitting parameter. The asymptote is again given by the value of *a* (Fig 4). This model is similar to the rational function (Appendix 1). It was found to outperform the Clench and negative exponential for both the training and testing datasets (Table 3).

**Figure 4.**
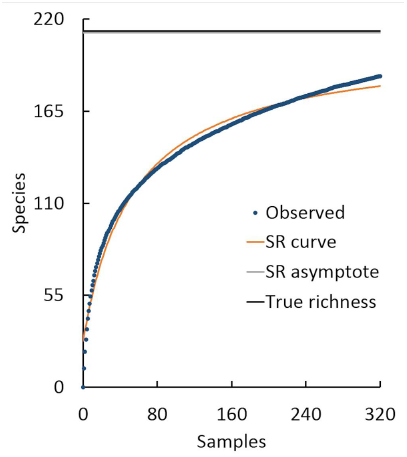
Accumulation curve for spider sampling in Gerês (Portugal). The result of searching for the best fitting asymptotic formula using symbolic regression is also shown.

**Table 3.** Comparison of three asymptotic equations used to estimate spider species richness in two forest sites. See Appendix 1 for formulas. Raw accuracy is the scaled mean squared error considering the entire observed accumulation curve (each formula was fitted to the curves using 4 to 320 samples) and weighted accuracy is this value weighted by the sampling effort at each point in the curve (where effort is the ratio between number of individuals and observed species richness). Note that lower values (in bold) are better as they reflect the deviation from a perfect estimator.

| Model | Raw accuracy | Weighted accuracy |
| --- | --- | --- |
| **Gerês (training)** |  |  |
| Observed | 0.113 | 0.037 |
| Clench | 0.055 | 0.018 |
| Negative exponential | 0.115 | 0.049 |
| Rational function | **0.045** | **0.012** |
| **Arrábida (testing)** |  |  |
| Observed | 0.103 | 0.031 |
| Clench | 0.038 | 0.010 |
| Negative exponential | 0.092 | 0.037 |
| Rational function | **0.032** | **0.008** |

### Developing the species-area relationship (SAR) and the general dynamic model of oceanic island biogeography (GDM)

For the Azorean spiders, the best fitting previous model (both highest r^2^ and lowest AICc) for the ISAR was the exponential model (Table 4). The SR run discovered roughly the same model, indicating, however, that the intercept (*c* term) was adding unnecessary complexity. A similar ranking of models was verified for bryophytes in the same region, revealing the robustness of the new model.

**Table 4.** Species area relationship (SAR) models for Azorean taxa and General Dynamic Models (GDM) of oceanic island biogeography for Canarian taxa. S = native species richness, A = area of the island and T = maximum time of emergence. Best models are indicated in bold.

| Model | Formula | r<sup>2</sup> | AICc |
| --- | --- | --- | --- |
| **SAR Azorean Spiders (training)** |  |  |  |
| Power | $S = 13.379 * A^{0.438}$ | 0.642 | 32.505 |
| Exponential | $S = 0.549 + 4.538 \log A$ | **0.780** | 28.102 |
| Linear | $S = 19.357 + 0.017A$ | 0.435 | 36.604 |
| Exponential (SR) | $S = 4.641 \log A$ | **0.780** | **23.319** |
| **SAR Azorean Bryophytes (testing)** |  |  |  |
| Power | $S = 181.625 * A^{0.803}$ | 0.666 | 78.085 |
| Exponential | $S = -27.824 + 57.114 \log A$ | **0.728** | 76.208 |
| Linear | $S = 196.215 + 0.259A$ | 0.617 | 79.295 |
| Exponential (SR) | $S = 51.889 \log A$ | 0.722 | **71.617** |
| **GDM Canarian Spiders (training)** |  |  |  |
| Whittaker | $S = -185.589 + 41.732 \log A + 17.776T - 1.022T^2$ | 0.873 | 110.350 |
| Fattorini | $\log S = 2.585 + 0.281 \log A + 0.157T - 0.009T^2$ | 0.941 | 105.025 |
| Steinbauer | $\log S = 3.367 + 0.098 \log A + 1.502 \log T - 0.454 \log T^2$ | 0.814 | 113.007 |
| SR | $S = 42.283 + 0.051A + 17.379T - T^2$ | **0.952** | **61.505** |
| **GDM Canarian Bryophytes (testing)** |  |  |  |
| Whittaker | $S = -176.599 + 66.602 \log A + 21.361T - 1.620T^2$ | **0.773** | **125.214** |
| Fattorini | $\log S = 4.544 + 0.137 \log A + 0.126T - 0.009T^2$ | **0.803** | **124.217** |
| Steinbauer | $\log S = 5.136 + 0.017 \log A + 1.063 \log T - 0.382 \log T^2$ | 0.612 | 128.963 |
| SR | $S = 192.660 + 0.075A + 20.702T - 1.576T^2$ | **0.785** | **124.841** |

For the Canary Islands, the best model for spiders was a linear function of area:

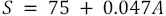

(r^2^ = 0.364, AICc = 65.631). Although it is easy to interpret, the explained variance is relatively low. The SR run reached a much higher explanatory power:

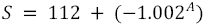

(r^2^ = 0.806, AICc = 57.320). In this case though, the equation is over-fitting to the few available data (7 data points), as this function is erratic creating a biologically indefensible model. The reason the ISAR is hard to model for the Canary Islands spiders is because we were missing the major component Time (Cardoso *et al.* 2010). This is depicted by the GDM, of which the best of the current equations was found to be the power model described by Fattorini (Fattorini 2009)(Table 4). Nevertheless, using SR we were able to find an improved, yet undescribed, model (Table 4). This represents a general model expanding the linear SAR:

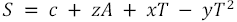

When tested with Canarian bryophytes, this new formulation is almost as good as the power model (Table 4).

## DISCUSSION

Symbolic regression has the advantage over most standard regression methods (e.g. GLM) in being fully flexible, allowing a much better fitting to data with similar interpretability to, for example, a linear regression. SR also has one or more advantages over other, commonly used, highly flexible regression (e.g. GAMs) or machine learning techniques (e.g. neural-networks): (1) numerical, ordinal and categorical variables are easily combined; (2) redundant variables are usually eliminated in the search process and only the most important are retained if anti-bloat measures (intended to reduce the complexity of equations) are used; (3) the evolved equations are human-readable and interpretable; and (4) solutions are easily applied to new data. Using SR we were able to “distil” free-form equations and models that not only consistently outperform but are more intelligible than the ones resulting from rigid methods such as GLM or “black-boxes” such as Maxent. We were also able to re-discover and refine equations for estimating species richness based on sampling curves and the ISAR and GDM from data alone.

All the examples presented in this work suggest that evolving free-form equations purely from data, often without prior human inference or hypotheses, may represent a yet unexplored but very powerful tool for ecologists and biogeographers, allowing the finding of hidden relationships in data and suggesting new ideas to formulate general theoretical principles.

### From particular relations to general principles

Scientific fields such as physics rarely rely on general statistical inference methods such as linear regression for hypothesis testing. The complexity of ecology made such methods an imperative in most cases. The method now presented not only allows the discovery of relationships specific to particular datasets, but also the finding of general models, globally applicable to multiple systems of particular nature, as we tried to exemplify. As mentioned, SR is designed to optimize both the form of the equations and the fitting parameters simultaneously. The fitting parameters usually are specific to each dataset, but the form may give clues towards some general principle (e.g. all archipelagos will follow an ISAR even if each archipelago will have its own c and z values). Although this aspect has not been explored in this study, we suggest two ways of finding general principles.

First, as was hinted by our estimators’ example, one may independently analyse multiple datasets from the same type of systems. From each dataset, one or multiple equations may arise. Many of these will be similar in form even if the fitting parameters are different. Terms repeated in several equations along the Pareto front or with different datasets tend to be meaningful (Schmidt & Lipson 2009). We may then try to fit the most promising forms to all datasets optimizing the fitting parameters to each dataset and look for which forms seem to have general value over all data.

Second, one may simultaneously analyse multiple datasets from the same type of systems but with a change to the general SR implementation. Instead of optimizing both form and fitting parameters, the algorithm may focus on finding the best form, with fitting parameters being optimized during the evaluation step of the evolution for each dataset independently. This parameter optimization could be done with standard methods such as quasi-newton or simplex (Nocedal & Wright 1999). To our knowledge, this approach has yet to be implemented, but it would allow finding general models and possibly principles, independently of the idiosyncrasies of each dataset.

### The need for human inference

Our results show that an automated discovery system can identify meaningful relationships in ecological data. Yet, as shown by our Canary Island spider SAR model, some equations might be very accurate but overfit the data. As with any relationship finding, either automated or human, correlation does not imply causation and spurious relationships are not only possible but probable given complex enough data.

Although the method here presented is automated, it is part of a collaborative human–machine effort. The possibility of exploiting artificial intelligence working together with human expertise can be traced back to Engelbart (Engelbart 1962), where the term “augmented intelligence” was coined to designate such collaboration. It has been subsequently developed and extended to teamwork involving one or more artificial intelligence agents together with one or (many) more humans, in diverse domains such as robotic teams (Yanco, Drury & Scholtz 2004) or collective intelligence for evolutionary multi-objective optimization (Cinalli 2015).

In ecological problems, human knowledge may play a fundamental role: 1) in the beginning of the process, when we must select input variables, building blocks and SR parameters; and 2) in the interpretation and validation of equations. The choice of equations along a machine-generated Pareto front should also take advantage of human expert knowledge to identify the most interesting models to explain the data. The researcher might then decide to disregard, accept or check equation validity using other methods.

### A priori knowledge

To some extent, it is possible to select *apriori* the type of models the algorithm will search for by selecting the appropriate variables and building blocks. Another way to take advantage of previous knowledge is to use as part of the initial population of equations some, possibly simpler, equations we know are related with the problem. For example, when searching for the GDM we could have given the algorithm multiple forms of the ISAR to seed the search process. This should be complemented with random equations to create the necessary variation for evolution.

### Fine-tuning the process

The number of options in SR is immense. Population size is positively correlated with variability of models and how well the search space is explored, but might considerably slow the search. Mutation rates are also positively correlated with variability, but rates that are too high might prevent the algorithm converging on the best models. The fitness measure depends on the specific problem and the type of noise expected.

The number of generations to let the search run is entirely dependent on the problem complexity and time available. Often the algorithm reaches some equation that makes immediate sense to the researcher and the process can be immediately stopped for further analysis of results. Sometimes several equations seem to make sense but are not entirely convincing, in which case several indicators can be used as a stop rule, such as high values of stability and maturity of the evolution process (Schmidt 2015).

The speed with which evolution occurs is extremely variable, depending on factors including the complexity of the relationships, having the appropriate variables and building blocks and the level of noise in the data. Fortunately, the process is easily adaptable to parallel computing, as many candidate functions can be evaluated simultaneously, allowing the use of multiple cores and even computer clusters to speed the search of equations.

### Caveats and alternatives

The SR approach is fully data-driven. This means it requires high-quality data if meaningful relationships are to be found. Also, it makes no a priori assumptions, so the final result might make no (obvious) sense, leading to spurious inferences, particularly if data are scarce or poor-quality, or if the right building blocks are not provided. Additionally, SR suffers from the same limitations of evolutionary algorithms in general. In many cases the algorithm may get stuck in local minima of the search space, requiring time (or even a restart with different parameters) to find the global minimum.

Many data mining techniques are regarded, and rightly so, as “black boxes”. SR is transparent in this regard, as variables are related through human-interpretable formulas. This is particularly important if the goal is to find equations with both predictive and explanatory power, building the bridge between finding the pattern and explaining the driving process, or if a general principle is to be suggested.

### The automation of science?

The methods here presented can be powerful additions to theoretical and experimental ecology, even if new conceptual hypotheses have to be created to accommodate the new equations. Such models could even be the only available means of investigating complex ecological systems when experiments are not feasible or datasets get too big/complex to model, using traditional statistical techniques.

This kind of techniques has led several authors to suggest the “automation of science” (King *et al.* 2009), where computers are able to advance hypotheses, test them and reach conclusions in largely unassisted processes. The SR potential as an exploratory step, to be reasoned alongside and proven with other methods is also exciting. The resulting formulas will help researchers to focus on initially imperceptible but interesting relationships within datasets and help guide the process of hypothesis creation.

## ACKNOWLEDGEMENTS

We thank Robert Whittaker, Stano Pekár, Michael Lavine and Otso Ovaskainen for comments on earlier versions of the manuscript; Carla Gomes and Ronan Le Bras for fruitful discussions around AI and Ecology. PAVB and FR were partly funded by the project FCT-PTDC/BIA-BIC/119255/2010 - “Biodiversity on oceanic islands: towards a unified theory”. LC was partially funded by UID/MULTI/04046/2013, from FCT/MCTES/PIDDAC, Portugal.

## DATA ACCESSIBILITY

All data available in Appendix 2.

## Appendix 1

Examples of general principles in ecology and of some of the 643 respective statistical models.

## Appendix 2

Data and settings used for all analyses in the paper (Eureqa file: http://www.nutonian.com/products/eureqa/).

## Appendix 3

Species, area and age for each Azorean or Canarian Island.

